# 36-year study reveals stability of a wild wheat population across microhabitats

**DOI:** 10.1101/2022.01.10.475641

**Authors:** Tal Dahan-Meir, Thomas James Ellis, Fabrizio Mafessoni, Hanan Sela, Ori Rudich, Jacob Manisterski, Naomi Avivi-Ragolsky, Amir Raz, Moshe Feldman, Yehoshua Anikster, Magnus Nordborg, Avraham A. Levy

**Affiliations:** Department of Plant and Environmental Sciences, Weizmann Institute of Science; Rehovot, Israel; Gregor Mendel Institute, Austrian Academy of Sciences, Vienna BioCenter; Vienna, Austria; Institute of Evolution, University of Haifa; Haifa, Israel; The Institute for Cereal Crops Improvement, Tel-Aviv University; Tel Aviv, Israel; Migal, Galilee Technology Center; Kiryat Shmona, Israel

## Abstract

Long-term genetic studies of wild populations are very scarce, but are essential for connecting ecological and population genetics models, and for understanding the dynamics of biodiversity. We present a study of a wild wheat population sampled over a 36-year period at high spatial resolution. We genotyped 832 individuals from regular sampling along transects during the course of the experiment. Genotypes were clustered into ecological microhabitats over scales of tens of metres, and this clustering was remarkably stable over the 36 generations of the study. Simulations show that it is difficult to explain this spatial and temporal stability using only limited dispersal, suggesting a role for fine-scale local adaptation to ecological parameters. Using a common-garden experiment, we showed that the genotypes found in distinct microhabitats differ phenotypically, further supporting the hypothesis of local adaptation. Our results provide a rare insight into the population genetics of a natural population over a long monitoring period.

## Introduction

The spatial and temporal scale of population genetic dynamics are key to understanding questions of local adaptation and the response to selection. For example, how does the scale of genetic variation correspond to that of environmental variation, and how quickly can populations respond to natural selection? To address these questions, we need to describe the spatial structure of genetic variation, and track how this changes over many generations.

Numerous studies have examined the spatio-temporal dynamics of wild plant populations over multiple years (Horovitz & Feldman 1991; Volis *et al*. 2004; Franks *et al*. 2007; Ozbek *et al*. 2007; Hendrick *et al*. 2016; Monnahan & Kelly 2017; Frachon *et al*. 2017; Troth *et al*. 2018; Fu *et al*. 2019; Todesco *et al*. 2020; Kolis *et al*. 2022). A few studies have examined the temporal scale of allele frequency change over decades, especially in long-lived vertebrates (Aguillon *et al*. 2017; Chen *et al*. 2019; Ashraf *et al*. 2021; Stoffel *et al*. 2021; Kelly 2022). However, there are essentially no genetic studies examining both spatial and temporal dynamics of wild plants’ populations over decades.

We examined the genetic diversity of a population of wild emmer wheat (*Triticum turgidum ssp. dicoccoides*) in Ammiad, Israel, which has been monitored for 36 years (Figure 1A). Wild emmer wheat is an annual, self-pollinating, allotetraploid cereal (genome BBAA). It is the progenitor of durum wheat (*T. turgidum ssp. durum*) and the maternal donor of the A and B sub-genomes of bread wheat (*T. aestivum*, genome BBAADD) (Levy & Feldman 2022). Its biodiversity is an invaluable resource for food security, due to the fact that major effect loci from wild emmer wheat can be transferred to the background of domesticated wheat. The Ammiad population shows high genetic diversity compared to other wild wheat populations (Levy & Feldman 1988; Felsenburg *et al*. 1991), together with variable topography (Figure 1, Movie S1) and a diversity of ecological niches within the scale of tens of metres (Anikster & Noy-Meir 1991; Noy-Meir *et al*. 1991b). The site is within a protected nature reserve, allowing repeated sampling of the population along transects that encompass considerable environmental variation. In this study we genotyped samples from nine harvests of these transects between 1984 and 2020, which allowed us to characterise the spatial and temporal dynamics of genetic change at Ammiad. Previous works suggesting that plants in Ammiad are adapted to microhabitats (Anikster *et al*. 1991; Noy-Meir *et al*. 1991b) were later disputed by Volis (Volis *et al*. 2015, 2016). Our dataset provides an almost unique insight into genetic change in a natural population of a predominantly selfing plant over time and space. This is especially relevant, because wild progenitors of modern crops such as wild emmer wheat serve as an immense reservoir of biotic and abiotic stress-resistance genes (Khoury *et al*. 2021) that could be utilised to improve commercial varieties. Sustained studies of progenitor species in their natural ecological niches can provide a better understanding of their survival in different ecological niches, and inform strategies for conservation of such resources *in-situ*, as well as in gene banks (Wintle *et al*. 2019).

**Figure 1.**
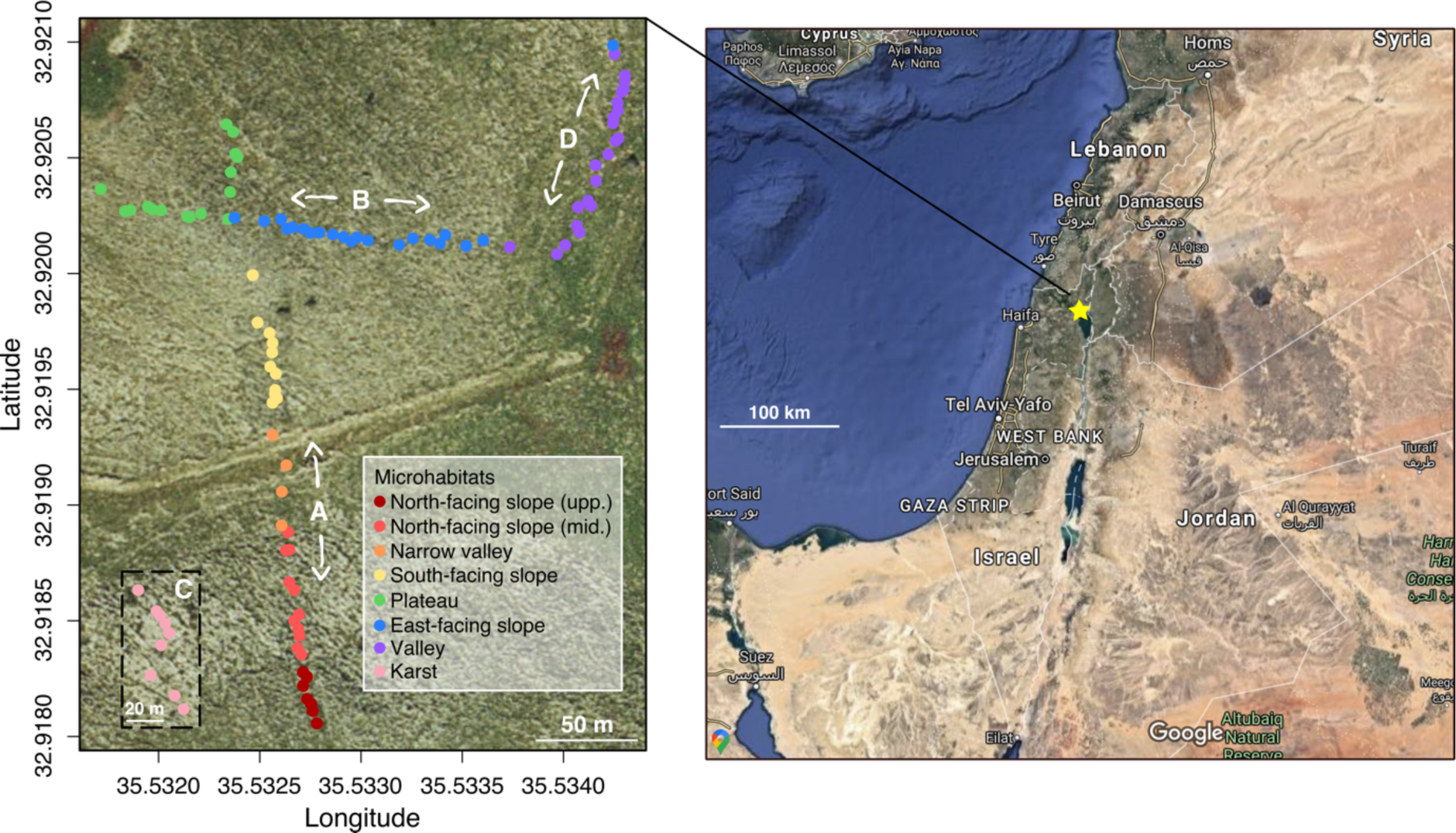
Ammiad wild-wheat population study. Location of Ammiad near the lake of Galilee (star, right and geographic coordinates, left). Sampling points of transects A, B, C, D collected in the years 1984, 1988, 1992, 1996, 2002, 2014, 2016, 2018 and 2020. Sampling points are shown on-site for transects A, B, D, but transect C, shown in the dashed rectangle, is located ∼500 metres north-west of A, B, D. Colours correspond to the different microhabitats labelled in the legend. The map on the left is taken from gov.map.co.il, and the map on the right is taken from Google Maps.

Here, we describe patterns of genetic variation at Ammiad in time and space. We then investigate whether an apparent association with microhabitat might be biologically meaningful by comparing the observed pattern of variation to an explicit model accounting for the limited seed and pollen dispersal of a predominantly self-fertilising plant. Finally, we discuss the implications of these results for local adaptation and conservation.

## Results

### The population is composed of groups of near-identical genotypes

We catalogued the genetic diversity at Ammiad by repeated sampling over 36 years. We marked 100 sampling locations with pegs and recorded GPS coordinates along four transects traversing seven ecologically distinct microhabitats (Anikster & Noy-Meir 1991); Figure 1, Dataset S1, Movie S1). We sampled seeds from the closest plant to each sampling location at each of nine time points, and sowed them in the greenhouse for genotyping. In total, we genotyped 832 individuals using a genotyping-by-sequencing (GBS) protocol (Poland *et al*. 2012), resulting in 5196 single copy, single-nucleotide-polymorphism (SNP) markers that were unambiguously positioned on the wild emmer wheat reference genome (Avni *et al*. 2017).

As expected for a highly selfing species (The 1001 Genomes Consortium 2016), the population is almost entirely composed of genetically distinct groups of nearly identical, highly homozygous individuals, clustered in space. Pairwise genetic identity follows a multimodal distribution that is characterised by pairs of plants that are either effectively identical, or share 70-90% of SNP alleles (Figure 2A). We unambiguously assigned all individuals to distinct genotype groups (DGGs) using a cut-off of >98% allele-sharing. This resulted in 118 DGGs over all nine time points, including 61 singletons (DGGs with a single member). The 10 largest DGGs encompass 65% of the samples, while the number of DGGs per year varied between 32 in 2014 to 49 in 1988 (Dataset S1). Rarefaction curves indicate that the sampling scheme was sufficient to detect almost all of the DGGs present close to the transects (Figure S1).

**Figure 2.**
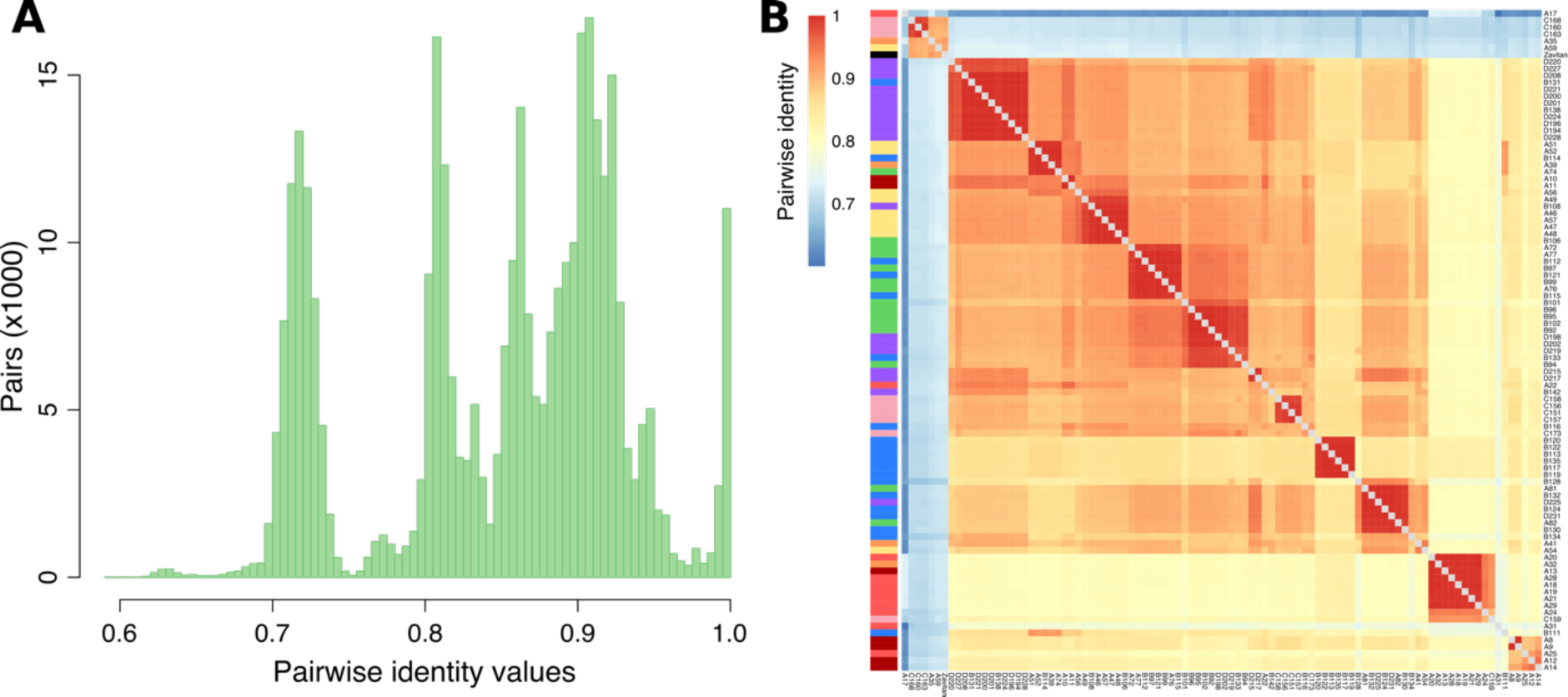
Ammiad population structure. (A) Distribution of pairwise identity values between the plants in the collection. (B) Population structure of Ammiad in the first year of collection (1984). Heatmap colours represent identity value for each pairwise comparison, annotation colours (on the left) correspond to the list of microhabitats in Figure 1.

We also identified 15 (1.8%) highly heterozygous plants, which is consistent with outcrossing in recent ancestors. Using SNPMatch (Pisupati *et al*. 2017), we were able to identify their parental DGGs (Figure S2B), suggesting that outcrossing happens between plants that are nearby. We estimated an average outcrossing rate of 3.7% across years in Ammiad, with detectable variation between the different habitats (1.8-8.8%, Table S1). These values are compatible with estimates in bread wheat (*Triticum aestivum*) (Enjalbert & David 2000).

### Genotypes are clustered by microhabitats and are stable through time

The spatio-temporal distribution of DGGs indicates two clear patterns (Figure 3). First, DDGs tend to cluster in space, which is expected for a self-compatible organism with limited dispersal. However, the clustering (Figure 3) and a principal component analysis of molecular variation (Figures S3A and S3B) show a striking concordance with the boundaries of ecological microhabitats, which were defined based on topographic and floristic data without knowledge of the genetic structure of the population (Noy-Meir *et al*. 1991b), Figure 3). For example, in transect A, there is a sharp transition between DGGs between the upper and middle north-facing slope across the 36 years of study (Figure 3A). Another example is the genotype at collection point A59 in a unique location under the shade of a *Ziziphus spina-christi* tree (Figure 3A). In contrast, in transect D, where there is negligible microhabitat differentiation, there is little spatial clustering of DGGs (Figure 3D). We found little evidence for a decay of relatedness that would be expected under a model of simple isolation by distance (Figure S3C and S4). Population differentiation measured by F_st_ was consistently higher between microhabitats than within, even when taking distance into account (Figures 4A, B), and genetic distances are strongly correlated with microhabitat. We also found greater differentiation among DGGs between microhabitats than would be expected by chance with the same level of spatial autocorrelation using a test based on Moran-spectral-randomization Mantel statistics (MSR-Mantel) that explicitly account for spatial distance (Figure S5). In summary, DGGs are associated with ecological microhabitats at the scale of tens of metres.

**Figure 3.**
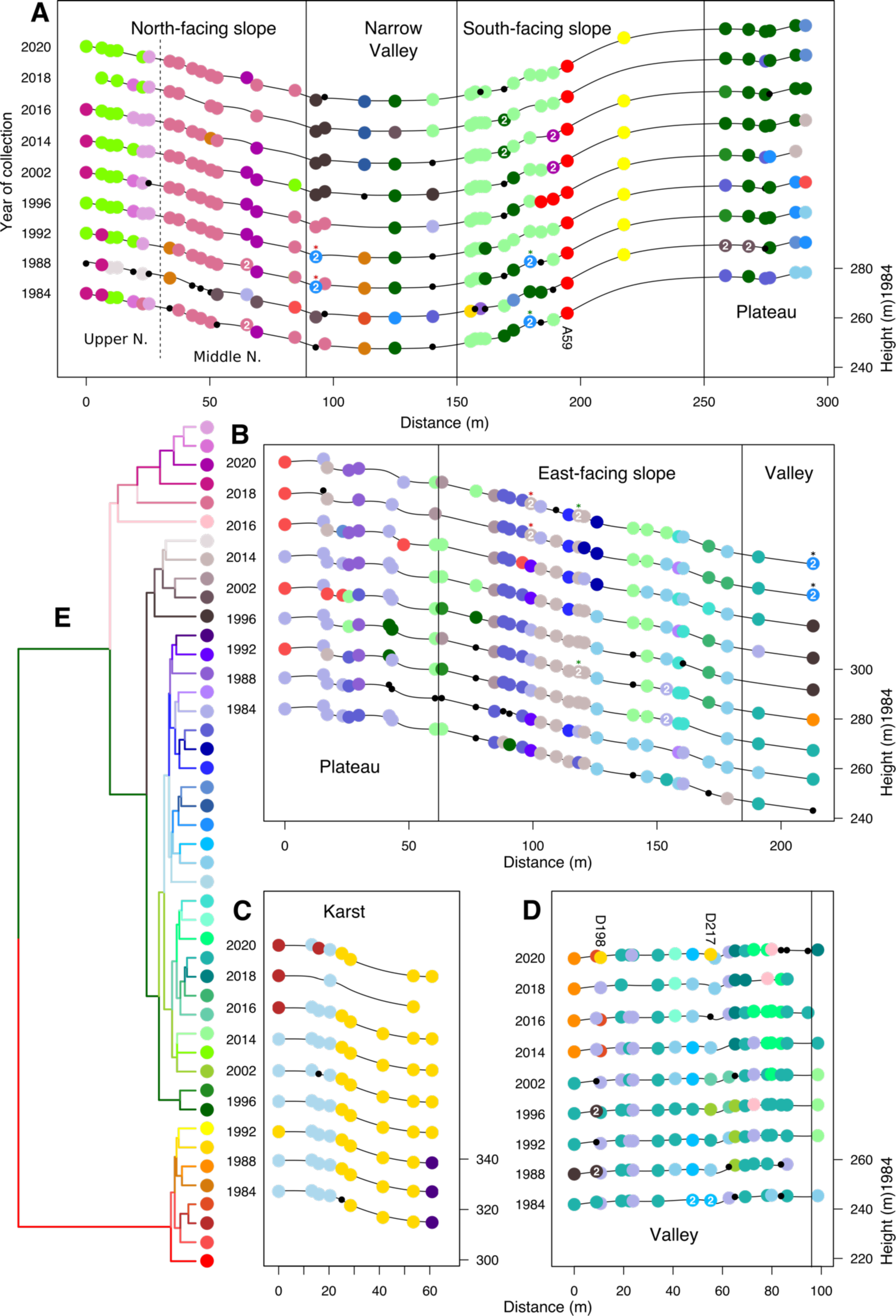
Spatio-temporal distribution of genotypes. (A-D) Distribution of samples coloured by distinct genotype group (DGG) in transects A, B, C and D. Y-axes indicate year of collection (left) and altitude for 1984 (right). Singleton samples are marked in small black dots, while doubleton groups are marked with “2” in the middle with the colour of their closest DGG. Doubleton groups that are closest to the same DGG were marked with coloured asterisks. (E) Dendrogram of DGGs with ≥ 3 samples, hierarchically clustered by UPGMA.

**Figure 4.**
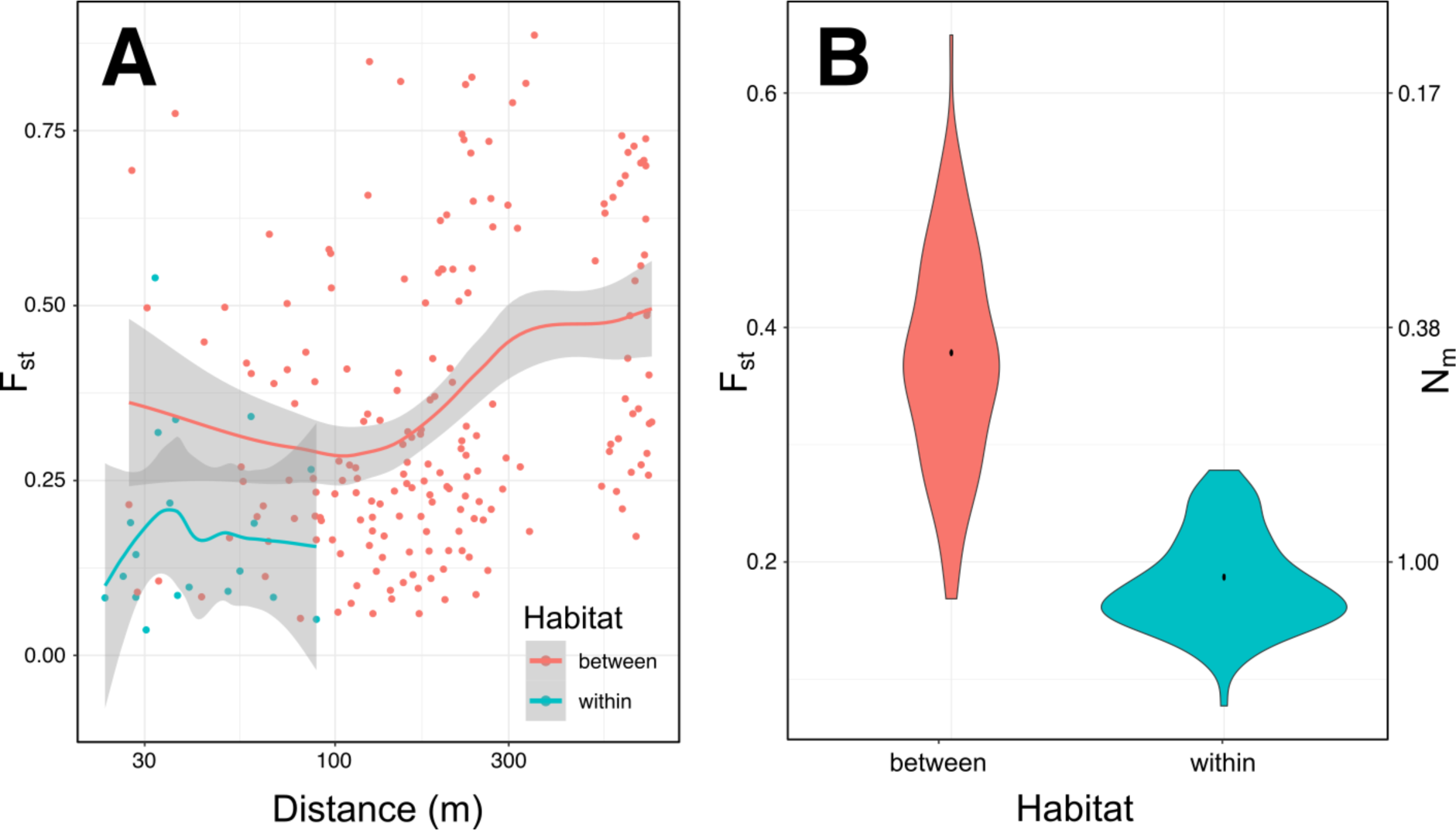
Spatial clustering at Ammiad. (A) Genetic differentiation between and within habitats. Plants within each habitat were grouped in 30m demes. Pairwise Fst was calculated among all demes, within and between habitats and plotted as a function of distance, measured as the average distance of plants in the two demes. Credibility intervals of a loess fit with degree 1 are shown in grey. Nm values (i.e., the theoretical number of migrants between parcels in an infinite island model) calculated are shown on the right axis. (B) Plants within each habitat were grouped in 30m demes. Pairwise Fst was calculated among all demes with average distance <50m between each other, within and between habitats. Average observed values are shown as a black dot. Uncertainty was estimated with 1000 bootstraps of the parcels considered for the comparisons. Average within-habitat Fst was higher than between-habitat Fst in 3% of the bootstraps. Nm values (i.e., the theoretical number of migrants between parcels in an infinite island model) calculated are shown on the right axis.

Second, the spatial clustering of DGGs was stable through time. For example, 33% of plants sampled in 1984 belonged to the same DGG as plants sampled at the same position in 2020. An analysis of molecular variance (AMOVA) showed that 28% of the genetic variation between samples is explained by differences between microhabitats, and that the habitat effect is highly significant (Table S2A, B and C). Conversely, time did not explain a significant proportion of variance (variance explained<1%), suggesting a strong stability of the genetic structure over time. These results suggest that space, and particularly microhabitats, might constrain the genetic structure of Ammiad to be stable over the scale of tens of generations.

### Neutral processes are unlikely to explain spatio-temporal stability

Our findings indicate that the genetic structure at Ammiad is more clustered into environmental microhabitats than would be expected under a null hypothesis of no clustering. However, it is clear that such a strict null hypothesis is not a good reference because it ignores the underlying biology of pollen and seed dispersal. To generate a more biologically realistic alternative hypothesis we tested whether the patterns at Ammiad can be explained using explicit demographic models that incorporate self-fertilisation and limited dispersal. The question is whether these neutral demographic processes - on their own - could generate the clustering into microhabitats observed at Ammiad.

We performed individual-based simulations of populations of plants that initially reflect the genetic structure at Ammiad, and subsequently evolve by seed dispersal and outcrossing only (see *Materials and Methods*). We simulated populations in a two-dimensional continuous space using the genetic structure at Ammiad in 1984 along one of those dimensions. We also recorded the microhabitat structure of those genotypes but did not model selection for those microhabitats. At each of 40 subsequent generations seeds dispersed, with some proportion of seeds being drawn from a two-year seed bank. In addition, some proportion of those seeds were assigned to be outcrossed and given a new genotype. At each generation we sampled a transect through the population and sample plants closest to 30 evenly spaced sampling points five metres apart. In this way we aimed to simulate a population evolving in the absence of selection and to sample it in a way that mimics the sampling of the study site at Ammiad.

We ran simulations using different values of outcrossing rate, plant density, seed dormancy and mean seed dispersal distance that approximate a range of biologically realistic values. Empirical values of outcrossing rates in wild wheat have been estimated to be as low as 0.5% (Golenberg 1988), although our estimates point to outcrossing rates of between 2% and 8% (mean = 3.7%; table S1). This species has a substantial seed bank with up to 30% of germinating seedlings coming from the previous year, but very few from previous years (Horovitz 1998). Average plant density in the Ammiad population was measured in 1984 as 4 plants/m^2^ (Noy-Meir *et al*. 1991a). We repeated this measurement in 2020 for transect A and found a similar density of 3.8 plants/m^2^ (Dataset S1). Seed dispersal occurs through a combination of gravity, ants, birds, grazing mammals and rodents, enabling both short and long-distance dispersal. We lack direct estimates of seed dispersal distances, but average gene flow, a composite of seed and pollen dispersal, was estimated to be 1.25 m per year (Golenberg 1987); since most dispersal is via seeds, the average seed dispersal distance is likely to be around 1 m per year. We aimed to simulate values above and below these estimates to allow for a range of biologically plausible values.

We compared the change in genetic differentiation among microhabitats in observed and simulated populations by tracking the change in F_st_ through time (Figure 5). At Ammiad F_st_ fluctuated between 0.08 in 1988 and 0.26 in 1996, but remained stable at around 0.15 on average throughout the period of study. When outcrossing rates in simulations were at their lowest value (0.5% per generation) F_st_ was similarly stable across time, especially when dispersal is also very limited (Figure 5, top row). However, for all other parameter combinations we observed a steady decrease in F_st_ in simulated populations over the 40 generations. The rate of decay was markedly faster as outcrossing rates increased, and depended surprisingly on dispersal distance, seed dormancy or plant density. This indicates that in the absence of selection, limited outcrossing of a few percent combined should be sufficient to erase any apparent sorting into microhabitats within the period of study examined here. Unless the outcrossing rate is much lower than we estimated, we cannot explain the observed microhabitat differentiation via neutral demographic processes.

**Figure 5.**
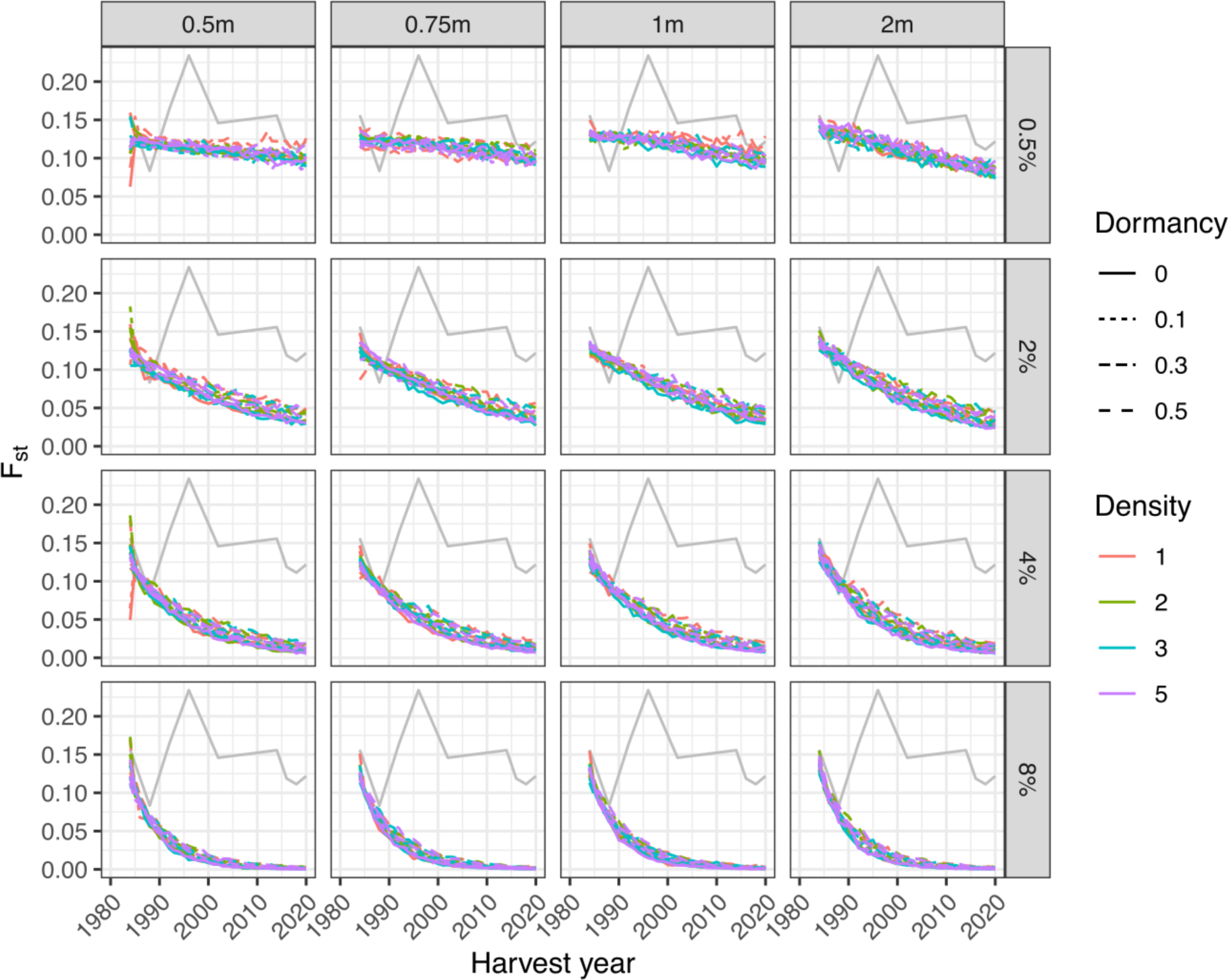
Genetic differentiation between microhabitats at Ammiad and in simulated populations. Grey lines show Fst between microhabitats across all transects at Ammiad. Coloured lines show average Fst over 100 replicate simulations that begin with the genetic structure at Ammiad in 1984 but evolve neutrally after that. Curves indicate different combinations of mean dispersal distance (grey boxes, panels left to right), outcrossing rate (grey boxes, panels top to bottom), proportion of seeds coming from the seed bank (dashed lines) and plant density per square metre (colours).

### Phenotypes measured in a common garden predict phenotypic differentiation between habitats

If there is a functional link between population structure and microhabitat differentiation, there is a prediction that habitats should be associated with differences in plant phenotypes. Owing to the challenging terrain and the status of the site as a protected conservation area it is not possible to estimate phenotypes with replication in a common-garden experiment under field conditions. We instead estimated phenotypes in a controlled net house experiment and used these data to predict phenotypic differentiation in the field.

To characterise the phenotypic variation between DGGs we grew six replicates of 95 DGGs in fully randomised blocks in a net house at the Weizmann Institute in conditions at Ammiad as closely as possible. We measured 14 phenotypes related to growth and reproduction (Figure S6) and estimated genetic values for each DGG using Bayesian generalised-linear models, accounting for differences between blocks. With the exceptions of germination date, number of spikes and number of tillers, broad-sense heritabilities (on the scale of the link functions; (de Villemereuil *et al*. 2016) were moderate to high (Figure 6A), indicating that substantial genetic variation exists for these traits.

**Figure 6.**
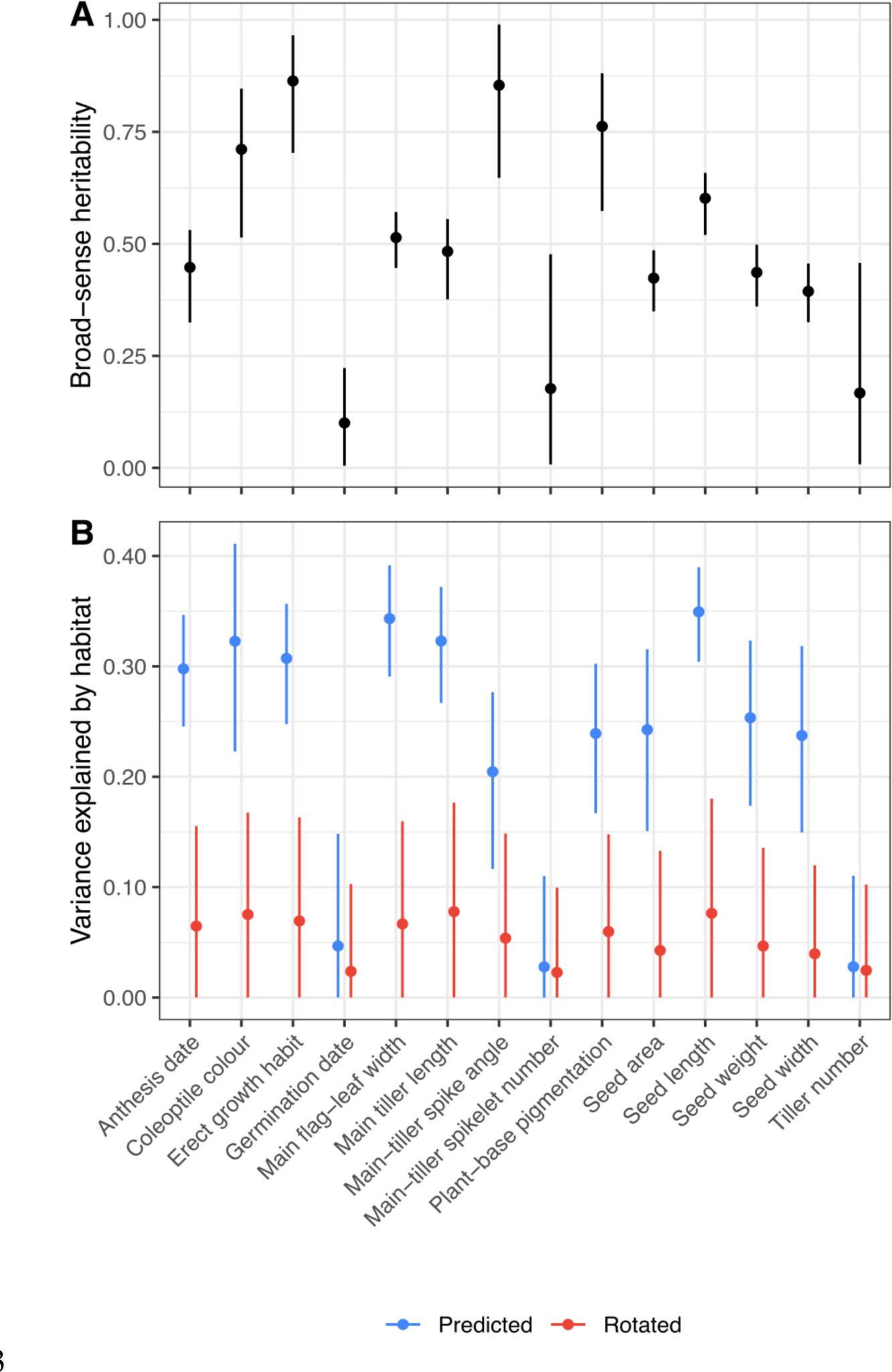
Heritability and variance explained by habitat for in predicted phenotypes at Ammiad. (A) Posterior mean and 95% credible intervals for broad-sense heritability of 14 phenotypes measured on six replicates each of 95 homozygous DGGs in a common-garden net house experiment. Because phenotypes display a range of error distributions, heritability is calculated on the scale of the corresponding link functions for comparison. (B) Plots show the posterior mean and 95% credible intervals for 801 field samples predicted (blue) from phenotypes measured in the net house experiment. Variance components are shown for the raw predictions based on real microhabitat positions, and for ‘rotated’ data (red) where microhabitat labels were offset by a random number of positions to break any correlation between phenotypes and microhabitats.

We then used phenotype estimates from the net house to predict phenotypic differentiation in the field. We assigned the 832 field samples a phenotype value for each trait from each of 1000 posterior draws of the corresponding linear models to generate 1000 sets of predicted phenotypes across time and space. A linear discriminant analysis that accounts for all traits jointly revealed clear clustering by microhabitat, with seed traits explaining the most separation (Figure S7; Table S3). For each trait separately, we then calculated the variance in each trait explained by habitat and year. Differences in habitat explained a very small proportion of the variation in the three traits that showed low heritability (tiller number, spikelet number and germination time; Figure 6B). For other traits, differences between habitats explained 19-33% of the phenotypic variation for all traits (based on posterior means; Figure 6B), whereas variation between years explained less than 10% of the overall phenotypic variance in the population. There thus seems to be differentiation in heritable traits between habitats which remains stable through time, consistent with the spatial and temporal patterns observed among DGGs.

We next asked whether spatial structure in phenotypes corresponds to the structure of microhabitats. We compared the predicted phenotypes to a ‘rotated’ dataset where the vector of phenotypes is offset by a random number of positions relative to the vector of habitat labels. This preserves the structure of phenotypes and habitat labels, but breaks any covariation between phenotype and habitat. In rotated datasets habitat typically explained less than 20% of the variation in phenotype. This is similar to that explained by the three predicted phenotypes with low heritability, but less than that explained by the eleven remaining phenotypes for more than 99% of predicted datasets (Figure 6B). Nevertheless, our results indicate that the genetic clustering by microhabitat is matched by phenotypic differentiation between microhabitats.

## Discussion

Studies of natural populations rarely have fine-scale geographic resolution or temporal resolution beyond the duration of a PhD or postdoc (Siepielski *et al*. 2009). Our study, spanning over 36 years of dense sampling, has both, and thus provides a unique dataset that could improve our understanding of gene flow and local adaptation on an ecological scale.

Our analysis revealed that the Ammiad population varies more in space than in time. We found unexpectedly strong long-term genetic stability, demonstrating that gene flow can effectively be highly limited (unless it is via pollen dispersal) in a wild cereal despite its wide distribution throughout the fertile crescent. Remarkably, the stable genetic clusters observed in Ammiad matched ecological microhabitats previously identified at the site, suggesting that both natural selection and limited dispersal have contributed to shaping population structure. Previous studies of wild populations have demonstrated local adaptation (Schemske 1984; Volis *et al*. 2015, 2016; Abebe *et al*. 2015; Sedlacek *et al*. 2016; Hendrick *et al*. 2016; Frachon *et al*. 2017; Gardiner *et al*. 2018) over large distances, but not at this spatial scale nor using long-term datasets. We also showed that the differences in genotypic compositions between habitats correspond to heritable differences in phenotypes, and that some of these phenotypes consistently differ among habitats. This further supports the hypothesis of local adaptation at the micro-environment level. Although we tried to replicate environmental conditions as closely as possibly, it is important to keep in mind that phenotypes were measured in a common garden rather than field conditions, and as such we cannot capture the precise environmental conditions at Ammiad, nor account for ways in which the phenotypes of individuals may vary when grown in different across microhabitats.

Our simulations showed that while the highly stable and structured population at Ammiad could be explained by very low outcrossing and seed migration, it is incompatible with the realistic parameters we estimated. This, together with the correlation between phenotypes-genotypes and microhabitats over a long experimental period, suggest that the population structure reflects adaptation to microhabitats, at least in part. The possibility that local adaptation occurs at such a fine spatial scale as in our study is intriguing and invites further experimental confirmation.

Although climate change was not yet an established fact in 1984, it became part of the motivation of Ammiad experiment. In retrospect, the increase in local ambient temperatures by ∼1.5 degrees Celsius (Figure S8A) and in atmospheric CO_2_ concentration by 70 ppm (Figure S8B) (Keeling *et al*. 2001) during the course of the experiment, could have led to selection that affected the population structure. However, only minor temporal changes were seen. This genetic temporal stability could be interpreted as the population not having had enough time to adapt due to the rapid pace of climatic changes, in which case we would expect a reduction in fitness of the population. However, plant density was similar in 1984 and in 2020, suggesting that, so far, this has not occurred. Wild emmer wheat might have evolved to be resilient to temperature fluctuations since its speciation 0.8 MYA ago (Levy & Feldman 2022) and that the recent climate changes might not have yet reached the threshold that endangers the species.

Our results are relevant to conservation strategies as well as to the assessment of biodiversity resilience under climate change. First, our study shows that *in-situ* conservation can be quite effective in preserving diversity, without the need to frequently sample seeds and store them in seed banks (*ex-situ*). Second, our results highlight that effective *in-situ* and *ex-situ* conservation strategies of wild plants biodiversity, particularly for wild grasses and crop progenitors such as wild emmer wheat, should consider habitat diversity as a potential reservoir of adaptive diversity. In fact, our long-term study would not have been possible if the site had not been declared a natural reserve for wild-wheat conservation by the Israel nature and parks authority. Indeed, this population is fenced and undergoes moderate grazing. Unrestricted grazing, or urbanisation, as seen in other parts of the Levant can reduce and sometimes eliminate wild grasses. The Ammiad reserve is relatively small (∼10 hectares) but contains much diversity. Establishing natural reserves of crop progenitors, partially protected from anthropogenic activities, does not necessarily require large areas and is a feasible goal that is critical for biodiversity conservation.

In conclusion, we provide a unique example of strong temporal stability, a counterpoint to recent studies demonstrating dramatic changes in local populations even between seasons (Bergland *et al*. 2014; Wittmann *et al*. 2017; Machado *et al*. 2021; Rudman *et al*. 2022). Fully explaining this stability will require further work, but, whatever the causes, such robustness in a rapidly changing world demonstrates the importance of conservation of wild relatives of crops in nature, as gene pools for breeding (Zsögön *et al*. 2021).

## Materials and Methods

### Sample collection and DNA extraction

Wild wheat spikes were collected in four different linear transects every 3 to 5 metres from sampling points marked with pegs and GPS coordinates (Dataset S1). Plants were assigned to a sampling point when collected in successive years in a radius of ∼ 1m from the peg. A single seed was selected for each plant and DNA was extracted from young seedlings. Tissue was collected and ground using mortar and pestle with the addition of Merck sea-sand in liquid nitrogen. DNA extraction was done using Macherey-Nagel NucleoSpin® Plant II extraction kit, according to the manufacturer’s protocol, eluting in 50 ul EB total. Concentrations and DNA quality were estimated using a NanoDrop® ND-1000 Spectrophotometer.

### Genotyping by sequencing (GBS) and variant calling

Plant genotyping was done according to the Genotyping-By-Sequencing method (Elshire *et al*. 2011; Poland *et al*. 2012), using *MspI* and *PstI-HF* restriction enzyme, and 96 barcodes of plate 384A (Poland *et al*. 2012). Samples were sequenced using Illumina NextSeq 550 mid-output 150 base-pairs single-end kits. A reference sample, Zavitan accession (Avni *et al*. 2017), was added to each run, undergoing the same plate GBS library preparation and sequencing. Samples were divided by barcodes using a python 2.7 code. Mapping to the WEW_v2.0 genome assembly (Zhu *et al*. 2019) was done using *bwa-mem (Li 2013)*. Conversion to binary alignment map (BAM) format, filtering for uniquely mapped reads with minimum QUAL score of 30, sorting and indexing was done using SAMtools (Li *et al*. 2009). Variant calling was done in parallel running on all 845 (832 Ammiad samples and 13 Zavitan controls) samples using GATK HaplotypeCaller 3.8 (Poplin *et al*. 2018). Filtering for quality (>30), read depth (>6), genotype quality (>15), maximum missing sites (10%) and no-indels was done using VCFtools (Danecek *et al*. 2011). Filtering for up to 10% heterozygous SNPs per plant was done using TASSEL5 (Bradbury *et al*. 2007). Filtered VCF was used for all further analyses (see Data and code availability).

### Pairwise genetic identity

We measured genetic identity using R (see Data and code availability) by calculating the pairwise identity values between all samples, divided by the number of comparisons. The pipeline also divides the samples into distinct genotype groups (DGGs) by setting a threshold of 98.1% identity of the variant sites, which is the lowest threshold where no sample is assigned to two or more DGGs, and constructs the dendrogram describing the relatedness between the DGGs. We performed hierarchical clustering R by UPGMA (method “average” in *hclust* function). All samples of the reference accession, Zavitan, were grouped together into one DGG, indicating that we can reliably group identical genotypes. The heatmap of samples from 1984 was done using *pheatmap* (https://cran.r-project.org/web/packages/pheatmap/index.html) with default hierarchical clustering.

### Temperature and CO_2_ concentration data

We obtained data on daily minimum and maximum temperatures from 1984 to 2020 from the Israeli Meteorological Service at the Israel government portal (https://ims.data.gov.il/he/ims/2). We obtained data on weekly atmospheric CO_2_ concentrations from The Scripps Institution of Oceanography UC San Diego (Keeling et al., 2001).

### Rarefaction, principal-component and identity=by-distance analyses

We performed a rarefaction analysis using R (see *Data and code availability*) by taking successively small subsamples of the DGGs present in each year, and fitting a polynomial curve to the number of unique DGGs sampled across subsamples. We repeated this 100 times for each year.

We performed principal component analysis on the VCF file of the entire Ammiad collection using *SNPRelate* R Package (Zheng *et al*. 2012). We calculated identity by distance for each transect separately. We calculated pairwise Euclidean distances using the *distHaversine* function from the *Geosphere* R package (Hijmans 2022). We performed Mantel tests with 1000 permutations to calculate the correlation between genetic identity and Euclidean distance matrices, using *mantel.test* in the *Ape* R package (Paradis *et al*. 2004).

### Outcrossing estimation and parental lines detection

To estimate outcrossing we calculated F_IS_ using the R package *hierfstat* (Goudet 2005). We calculated confidence intervals as the 2.5% and 97.5% quantiles of 100 bootstraps using the function boot.ppfis. We estimated outcrossing rates (*f*) using the relationship *F_IS_ = (1-f)/(1+f) (Wright 1969)*. To detect the parental DGGs of heterozygous plants we used SNPMatch (Pisupati *et al*. 2017) by comparing each plant to a database of all DGGs.

### Microhabitat clustering analysis

We tested whether population differentiation, measured by F_st_, is higher when measured between plants occurring in different microhabitats than in the same microhabitat. To this goal we subdivided plants into demes with a maximum distance between plants of 30m. We then computed the pairwise F_st_ between demes using the R package *pegas (Paradis 2010)*. To correct for the effects of isolation by distance and spatial structure we stratified comparisons by distance, measured as the distance between the average locations of plants in the two demes.

To account for spatial autocorrelation we first performed a partial Mantel test using the function *multi.mantel* from the R package *phytools (Revell 2012)*. Specifically, we considered as a response variable the pairwise genetic distances used to define DGGs. We tested for the effect of habitat, represented as a distance matrix with value “0” for plants found in the same microhabitat and “1” for plants found in two different microhabitats. In addition, we included in the test a matrices of spatial and temporal distance We estimated spatial distances based on latitude and longitude using the function *earth.dist* from the R package *fossil* (https://palaeo-electronica.org/2011_1/238/index.html). We detected a significant effect of space and microhabitat (p-values <0.005 for both matrices) and a non-significant effect of time (p-value = 0.863).

Mantel and partial Mantel tests have been severely criticised for their inaccuracy in presence of high spatial autocorrelation (Legendre *et al*. 2015; Crabot *et al*. 2019). To account for these known biases, we used the approach proposed by (Crabot *et al*. 2019), which uses a Moran spectral randomization procedure to generate random samples which maintain the same autocorrelation properties of the original dataset. To this goal, we used the R packages *sdpep* to process the spatial matrices and the *msr* function of the package *adespatial (Guénard & Legendre 2022)*. The distances above were transformed into Euclidean distances with function *cailliez* from the R package *adegenet (Jombart & Ahmed 2011)*. Using this approach, we tested the effect of habitat on both DGGs and pairwise genetic distances, using 1000 randomizations. In both cases the effect of microhabitat was significant (p-value<0.05).

We computed an analysis of molecular variance using the R package *pegas (Paradis 2010)*, using plants sampled across all years. We included terms for microhabitat and year of collection (nested within microhabitat) in the model.

### Individual-based simulations

To investigate whether neutral demographic processes could maintain the degree of clustering observed at Ammiad in 1984 over the period of study we performed individual-based simulations of populations of plants evolving via seed dispersal and outcrossing only. An R package, *simmiad*, is provided to perform these simulations (DOI:10.5281/zenodo.8116105).

We simulate populations of *N* plants evolving in a uniform two-dimensional habitat beginning with a structure that mimics that found at Ammiad with densities of 1, 2, 3 or 5 plants/m^2^. One challenge is that the real population is sampled along a one-dimensional transect, but simulations need to be seeded in two dimensions. To do this we first concatenate the vector of genotypes from all transects in 1984, arrange them along sampling points five metres apart along one axis of the habitat, then copied this structure in the perpendicular direction to create bands of identical genotypes in a grid. So as not to bias the measurement of the structure to any single region of the populations we ‘rotated’ the concatenated transect by a random number of positions in each simulation replicate. Sampling points are also assigned the same microhabitat label as they had at Ammiad, but we do not model selection for microhabitat. To ensure the population is at the correct density we then draw a sample with replacement of size *N* of plants on the grid to generate a population at generation 1.

At 40 subsequent generations, *N* plants are sampled with replication to produce *N* seeds. Wild emmer wheat has a two-year seed bank, so from generation 3 onwards, seeds are drawn from the previous generation with probability 1-*p*, and from two generations previous with probability *p,* for values of *p* of 0.5, 0.3, 0.1 or 0. Seeds are randomly assigned as outcrossed with probabilities 0.25%, 0.5%, 1%, or 2%. Since we have no information about seed dispersal distance and are only concerned with tracking unique genotypes, outcrossed seeds are assigned a new unique genotype, just as outcrossed seeds in the real sample would appear as a unique DGG. Offspring disperse from the mother in a random direction at a distance drawn from an exponential distribution with a mean of 0.5, 0.75, 1 or 2m. To eliminate edge effects, the habitat is modelled as a torus, such that if dispersal takes a seed over the edge of the habitat it is reflected back to the opposite side. At each generation we then draw a transect of 30 sampling points 5 metres apart through the middle of the population, and sampled the plant closest to each sampling point, replicating the sampling procedure in the field.

We measure microhabitat differentiation on each sampled transect as

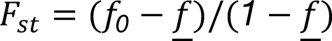

where *f_0_*is the is the probability that two individuals in the same subpopulation are the same DGG, and <u>*f*</u> is the probability that two individuals from the total population are the same DGG. For each combination of plant densities, outcrossing rates, seed bank parameter *p*, and seed dispersal distances we performed 100 replicate simulations.

### Common-garden experiment

We established a common garden experiment in a net-house in the Weizmann Institute of Science. We grew plants from 95 distinct homozygous DGGs from DGGs represented by at least two samples, as well as the reference genotype ‘Zavitan’, in six fully randomised blocks in a net house. In order to keep conditions as similar as possible to the conditions in Ammiad site, we sowed seeds in mid-November in Terra Rossa soil collected at Ammiad. Plants were subject to natural winter rainfall, and we provided additional irrigation on two occasions when plants were close to desiccation.

We measured 4 phenotypes related to vegetative growth: (germination date; presence/absence of coleoptile anthocyanin pigmentation; erect growth habit, scored from 1 to 5; plant-base pigmentation during tillering, scored from 1 to 3), six phenotypes related to anthesis (anthesis date; main tiller length (cm) from the soil to the base of the spike; log main-tiller flag-leaf width (cm); main tiller number of spikelets; main tiller spike angle, scored from 1 to 3; number of tillers) After seed collection we measured four additional phenotypes based on 10 dry seeds per plant: seed area (mm^2^), seed length (mm), seed width (mm) and the weight of 10 seeds (grams) using VIBE QM3 seed analyser. Raw phenotypic data are provided in Dataset S2.

We estimated the posterior distribution of genetic values for each trait using a generalised linear mixed model in the R package *brms (Bürkner 2017)*. We model the phenotype *z_ijk_* with link function *f(z)* (with inverse function *f‘*(*z*)) of a plant *i* of genotype *j* in block *k* as

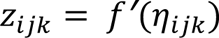

where

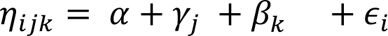

*α* is an intercept, *γ*_$_ is the mean phenotype of DGG *j*, β_%&_ is the mean phenotype for experimental block *k*, ε_#_ is a residual error term (for phenotypes with normally-distributed errors), and:

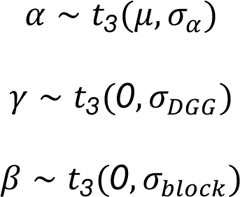

We used the default priors in *brms* for the mean and variance components. The 14 traits measured were a mixture of normally-distributed, Poisson, binary and ordinal variables, so we used identity, log, logit, and cumulative-logit link functions respectively, as appropriate. We standardised normally-distributed traits by mean-centering and dividing by the standard deviation. We calculated broad-sense heritability on the scale of the link function following (de Villemereuil *et al*. 2016) as

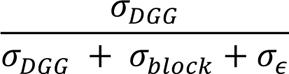

Note that for models with non-identity link functions σ_ε_ is only defined on the scale of the data, and does not contribute to heritability estimates on the scale of the link function (McCullagh & Nelder 1989; de Villemereuil *et al*. 2016).

We used the linear models to predict phenotypes in the field. We drew 1000 values from the posterior distribution of each phenotype for each of the 832 samples in the field data set (excluding 117 empty sampling plots) using the *brms* function ‘posterior_predict’. This function simulates phenotype values based on the genetic values for each DGG plus inter-individual error, accounting for uncertainty in both. For the 26 samples that were not included in the net-house experiment, we used *brms* to draw a phenotype value from the posterior distribution of genetic values of phenotyped DGGs. Since these values vary across posterior draws this allows us to assign phenotypes to the missing genotypes without biassing the distribution of phenotypes, and renders tests of association with microhabitat conservative.

We performed a linear discriminant analysis of the predicted phenotypes in the field as a function of microhabitat using the *MASS* library in R (Venables & Ripley 2002). For this analysis we used posterior mean phenotypes for each DGG, using the 806 plants from DGGs that were included in the net-house experiment.

We then estimated the variance in each phenotype explained by habitat and year in each predicted dataset using *lme4* (Bates *et al*. 2015). To generate a null distribution for the association between phenotype and habitats we used a ‘rotation’ approach similar to that used to test sorting by microhabitat. We concatenated transects within each year into a single circular transect and rotated the vector of predicted phenotypes by a random number of positions. This preserves spatial structure in phenotypes and microhabitats but breaks any association between them. We performed a single rotation for each of 1000 realisations of phenotype predictions. For each realisation we recalculated the variance in rotated phenotype explained by microhabitat and year, and compared the distribution of these variance estimates to those for the corresponding predicted dataset.

## Acknowledgements

We thank T. Azulay, T. Grinevich, C. Melamed-Bessudo, O. Gross, R. Epstein, Z. Meir and members from the Institute for Cereal Crops Improvement at Tel-Aviv University for plant material collection. We thank A. Distelfeld and members of the Institute of Evolution, University of Haifa, for using their VIBE QR3 seed analyzer, and J. Müller for the drone movie. We also thank N. Barkai, Y. Eshed, D. Filiault, K. Swarts, S. Nowoshilow, N. De-Malach, R. Pisupati, R. Burns, Y. Voichek, P. Clauw, E. Kosman and the Levy and Nordborg labs for fruitful discussions.

## Data and code availability

Sequencing data generated in this study are available in the National Center for Biotechnology Information (NCBI) Sequence Read Archive (SRA) under the accession number PRJNA779576. All data processing and data analysis are publicly available on GitHub (https://github.com/TalDM/Ammiad) and Zenodo (DOI:10.5281/zenodo.8116655). An R package, *simmiad*, is provided to perform the individual-based simulations (DOI:10.5281/zenodo.8116105).

## Author Contributions

T.D-M., M.F., Y.A. and A.A.L, conceived and designed the study. T.D-M. carried out the molecular work, SNP calling, genetic and phenotypic analyses with input from T.J.E., F.M., M.N. and A.A.L. T.J.E. performed the simulations and phenotypic analyses with input from T.D-M., M.N. and A.A.L. F.M. carried out the population differentiation analysis, with input from T.D-M. and A.A.L. T.D-M., O.R. and A.A.L, carried out the common-garden experiment. T.D-M., H.S., J.M., N.A-R., A.R., M.F., Y.A, and A.A.L carried out the collection. T.D-M., T.J.E., F.M, M.N. and A.A.L wrote the manuscript. M.N. and A.A.L supervised the project.

## Supplementary Files

**Movie S1 (separate file).** Drone video of the different transects and microhabitats in Ammiad wild wheat reserve.

**Dataset S1 (separate file).** Dataset of Ammiad samples by genotype, coordinates and spike density measured at transect A in 2020.

**Dataset S2 (separate file).** Dataset of plant and seed phenotypes measured in the common garden experiment.

---

**Figure S1.**
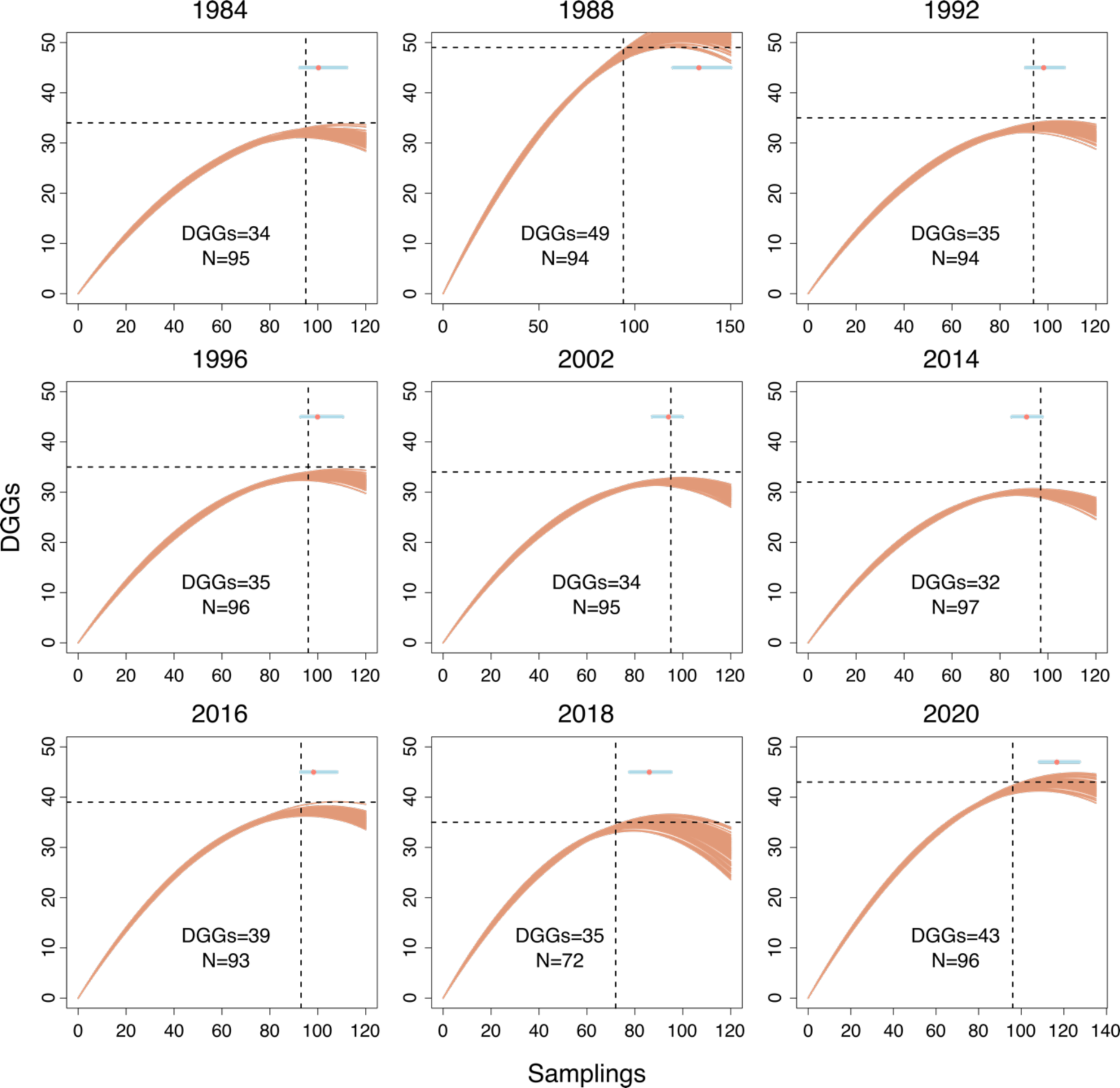
Rarefaction curves. Adjusted 2^nd^ degree polynomials of 100x backwards sample of unique genotype number and number of samples collected per year. Dashed horizontal line represents the observed number of unique genotypes per year, dashed vertical lines represent number of samples per year. Light blue line and red dot represent range and mean of inflection points over replicate subsamples respectively.

**Figure S2.**
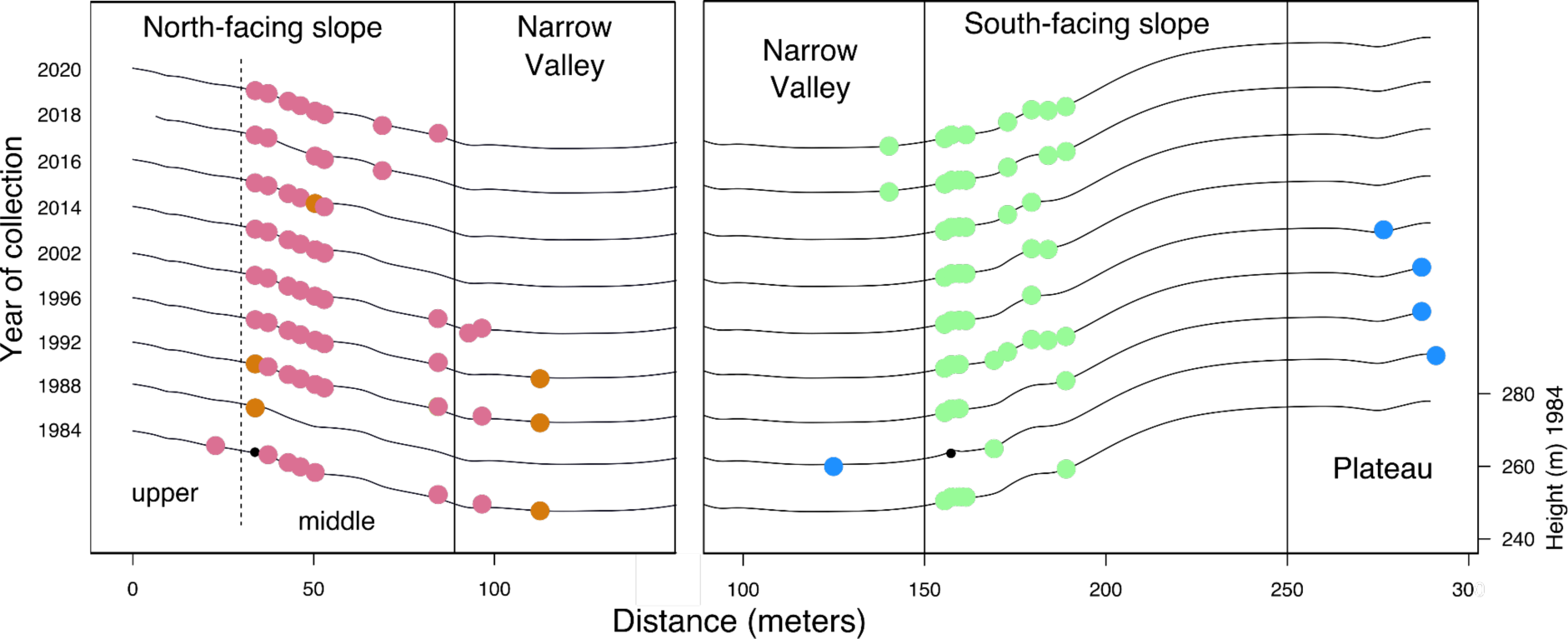
Outcrossing in Ammiad. (A) Outcrossing over time in Ammiad. The red continuous line represents the estimated average decrease of 0.06% every year, with confidence intervals shown as dashed lines. The highest rate of outcrossing is estimated for the middle north-facing slope, consistently with the identification of two highly heterozygous plants in this habitat from 1984 and 1988 (A17 and A20, respectively). Individual yearly estimates are shown as black dots. (B) Parental DGGs of 1984A17 (left panel) and 1988A47 (right panel) detected using SNPMatch. F1 plants are marked as a black dot.

**Figure S3.**
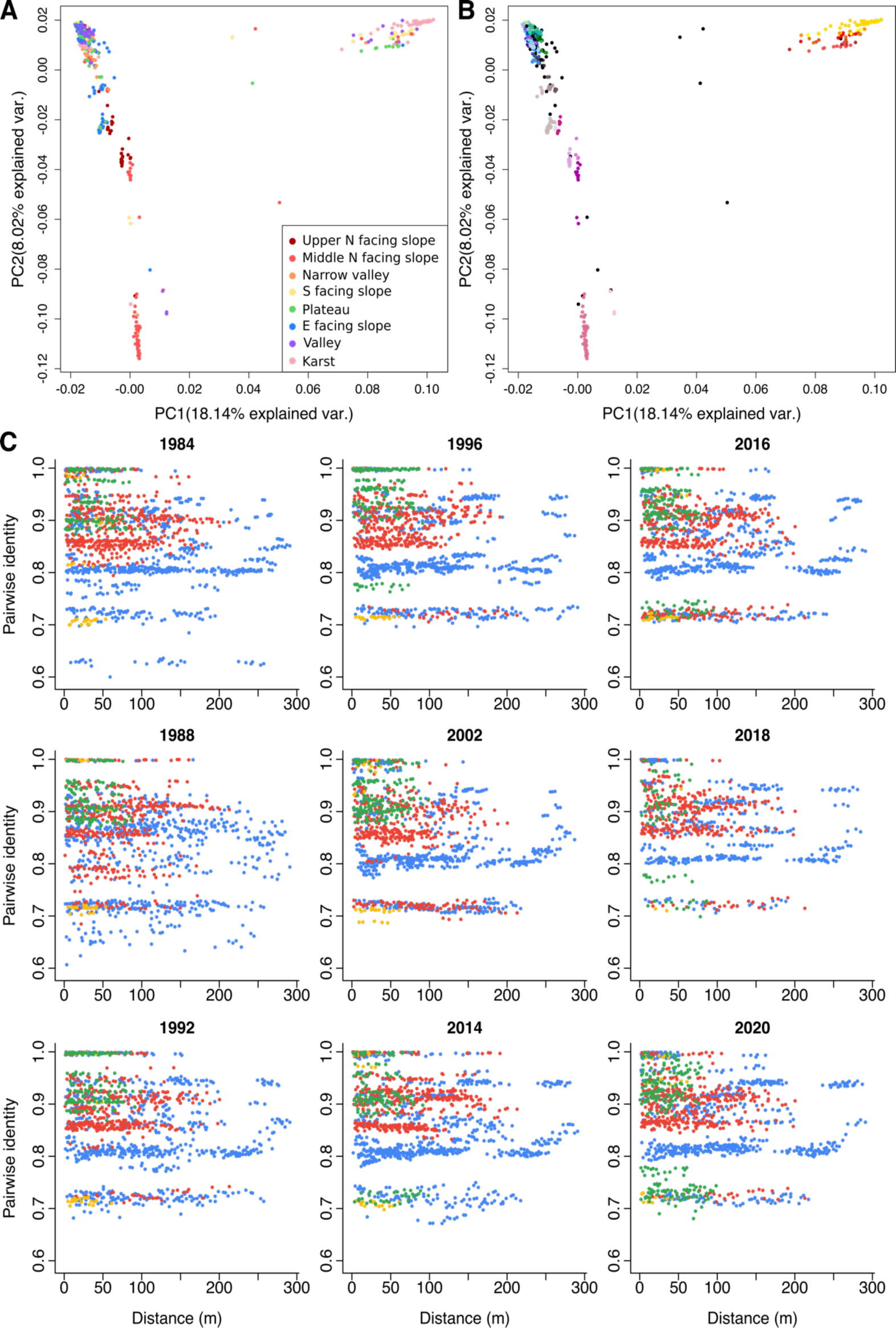
Principal Component Analysis (PCA) and isolation by distance of Ammiad collection. (A) PCA coloured by microhabitat. (B) PCA coloured by DGGs, black dots represent singleton DGGs. (B) Scatter plots of pairwise genetic identity and Euclidean distance (metres). Subplots show years of collection, and colours represent the different transects (A=blue, B=red, C=yellow, D=green).

**Figure S4.**
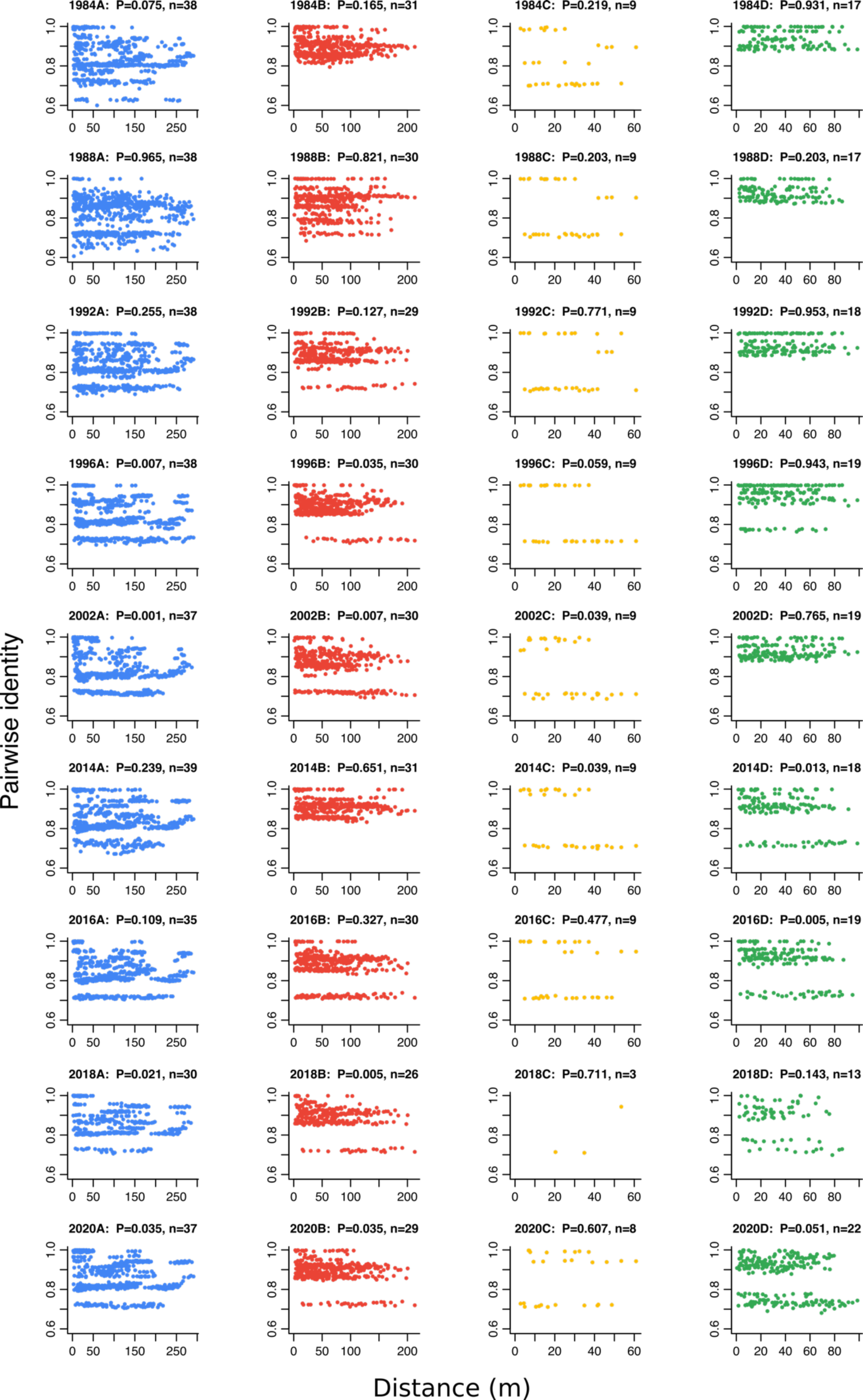
Correlation and significance of isolation by distance in Ammiad collection. Pairwise genetic identity by Euclidean distance (metres). Subplots show years of collection (rows) and transects (columns). The title of each plot indicates its number of samples and p-value from a Mantel test with 1000 permutations.

**Figure S5.**
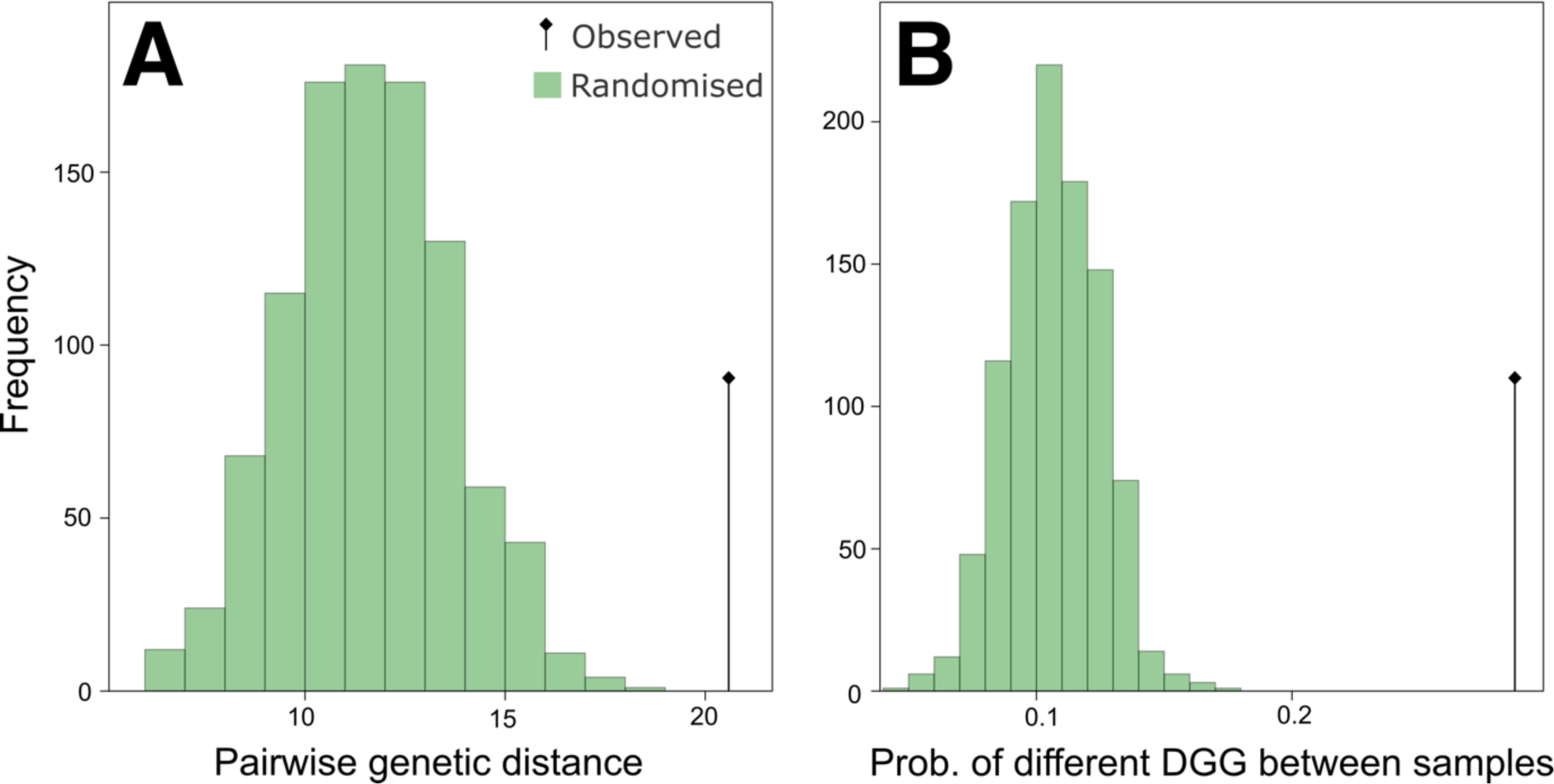
Moran Spectral Randomization for Mantel test to test the effect of habitat on genetic structure accounting for spatial autocorrelation. The histograms represent 1000 spatially constrained randomizations of Mantel statistics (MSR-Mantel). Each randomization was constrained to have a spatial autocorrelation comparable to the observed data, controlling for dispersal and other random factors. For each randomization the effect of habitat on pairwise genetic distances (A) and the probability of two samples to belong to the same DGG (B) was measured, and compared to the observed (black line and dot), in order to, test for the role of habitats in determining the spatial autocorrelation in genetic similarity in Ammiad. For both A and B, we observed a significant role of habitat in shaping the autocorrelation and the genetic distances in Ammiad, both with p-value <0.01.

**Figure S6.**
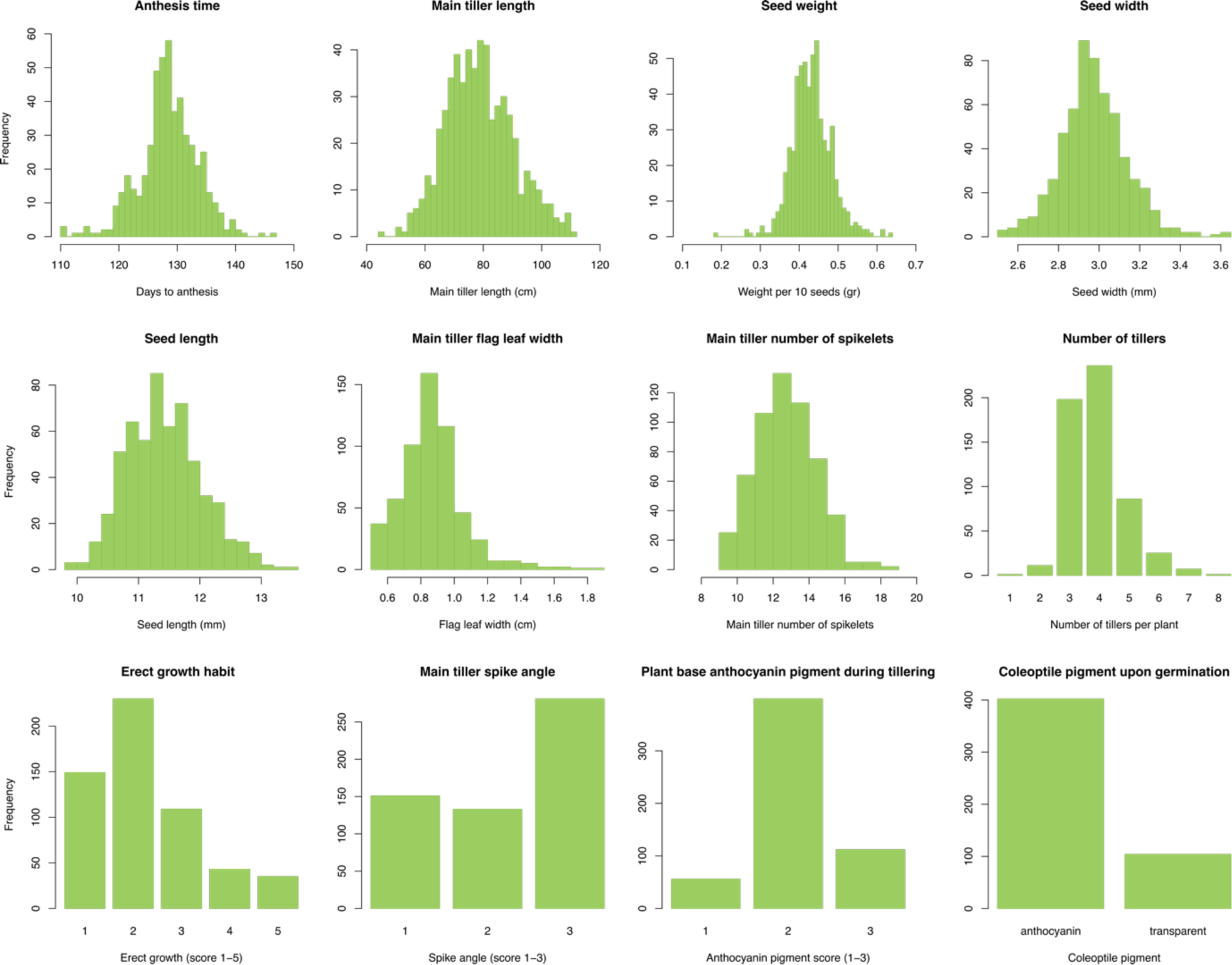
Distribution of phenotypes measured in the common-garden experiment.

**Figure S7.**
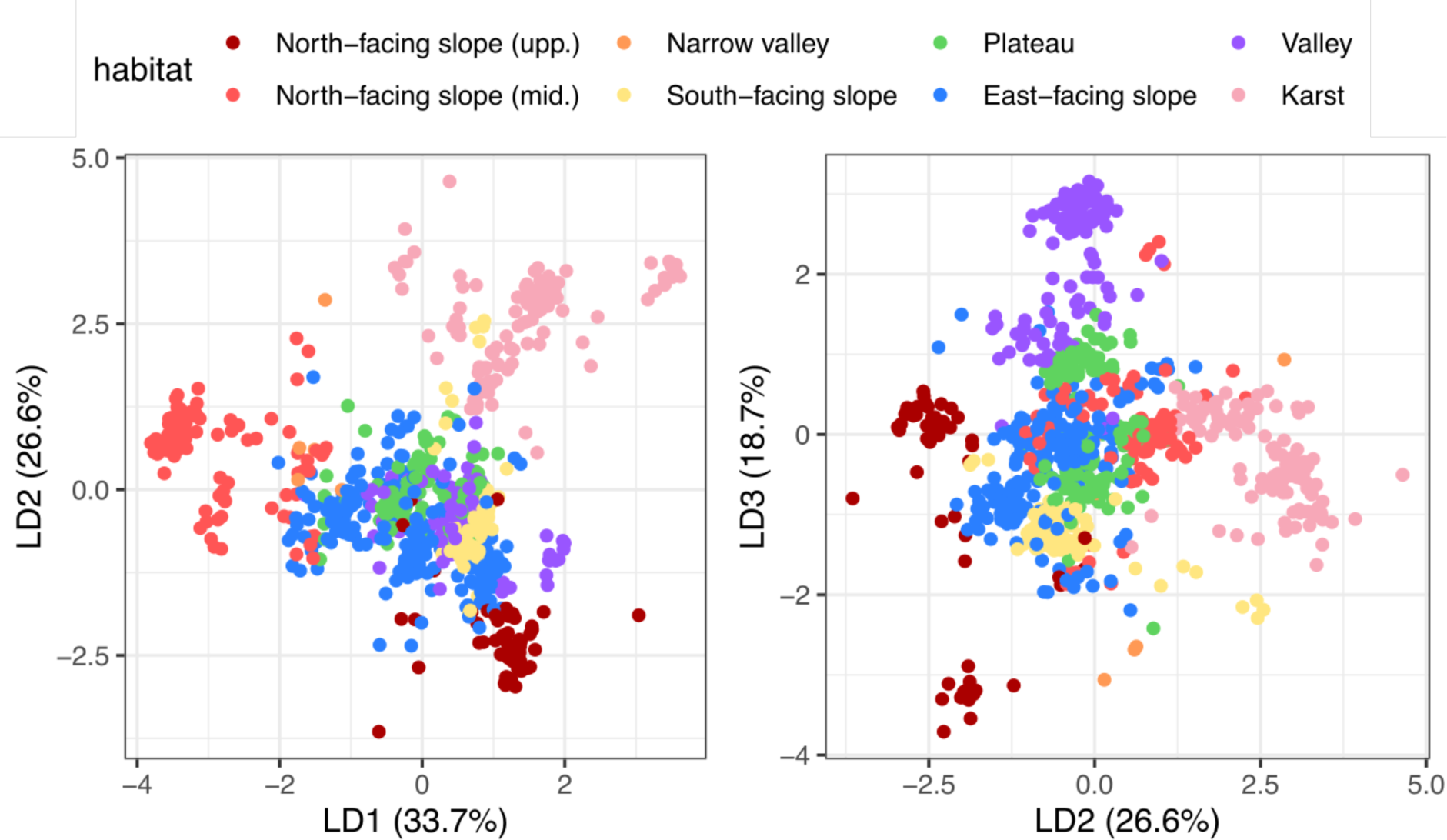
Phenotype-habitat linear discriminant analysis. Projection of the first three dimensions of the linear discriminant analysis of the posterior-mean predicted phenotypes of 801 samples from the wild wheat Ammiad population.

**Figure S8:**
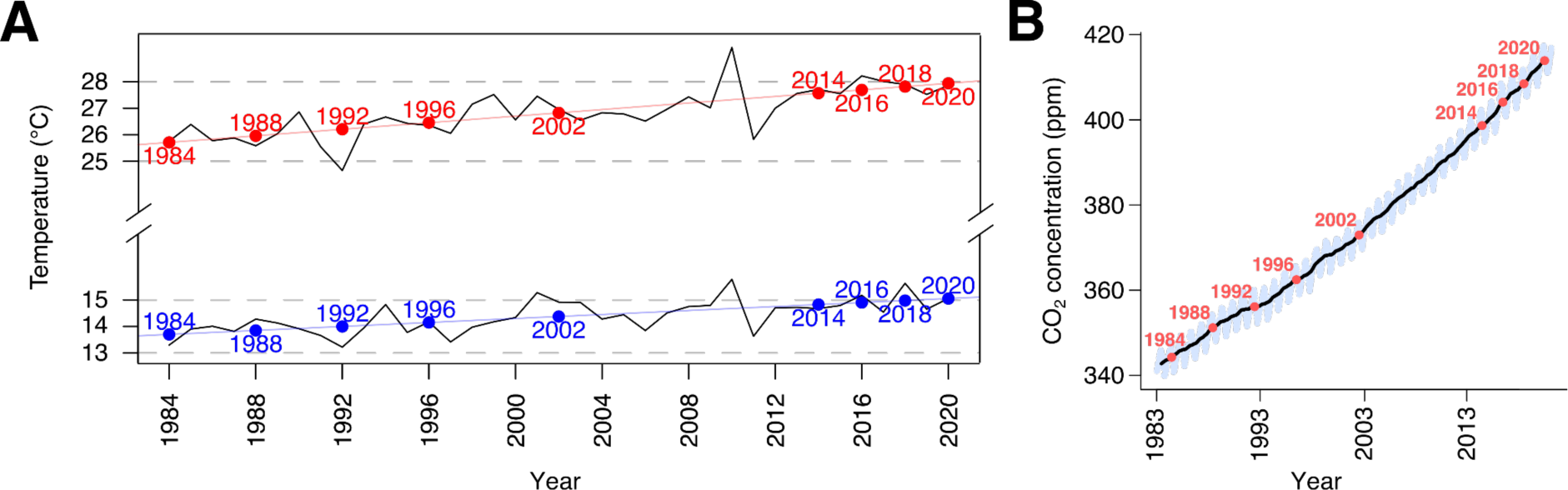
Climatic changes during the course of the Ammiad experiment. (A) Average yearly minimum (blue) and maximum (red) temperature measured in Ayelet Haschahar weather station from 1984 to 2020. Collection years are marked on the respective linear model regression line. (B) CO2 concentration 1983 to 2021. Data obtained from the Mauna Loa record, Scripps Institution of Oceanography, UC San Diego.

**Table S1.**
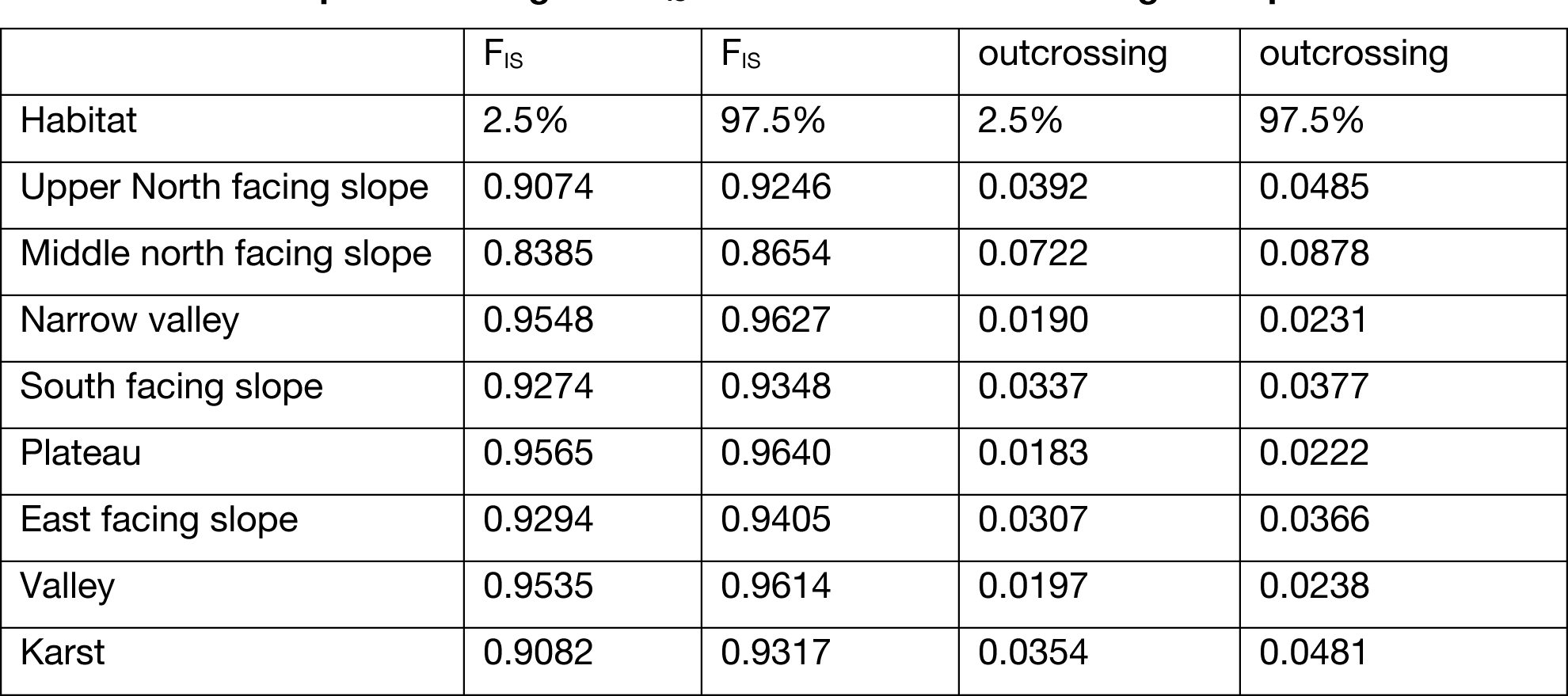
Bootstrap-based ranges of FIS and estimated outcrossing rates per habitat.

| | $F_{IS}$ | $F_{IS}$ | outcrossing | outcrossing |
| --- | --- | --- | --- | --- |
| Habitat | 2.5% | 97.5% | 2.5% | 97.5% |
| Upper North facing slope | 0.9074 | 0.9246 | 0.0392 | 0.0485 |
| Middle north facing slope | 0.8385 | 0.8654 | 0.0722 | 0.0878 |
| Narrow valley | 0.9548 | 0.9627 | 0.0190 | 0.0231 |
| South facing slope | 0.9274 | 0.9348 | 0.0337 | 0.0377 |
| Plateau | 0.9565 | 0.9640 | 0.0183 | 0.0222 |
| East facing slope | 0.9294 | 0.9405 | 0.0307 | 0.0366 |
| Valley | 0.9535 | 0.9614 | 0.0197 | 0.0238 |
| Karst | 0.9082 | 0.9317 | 0.0354 | 0.0481 |

**Table S2A.**
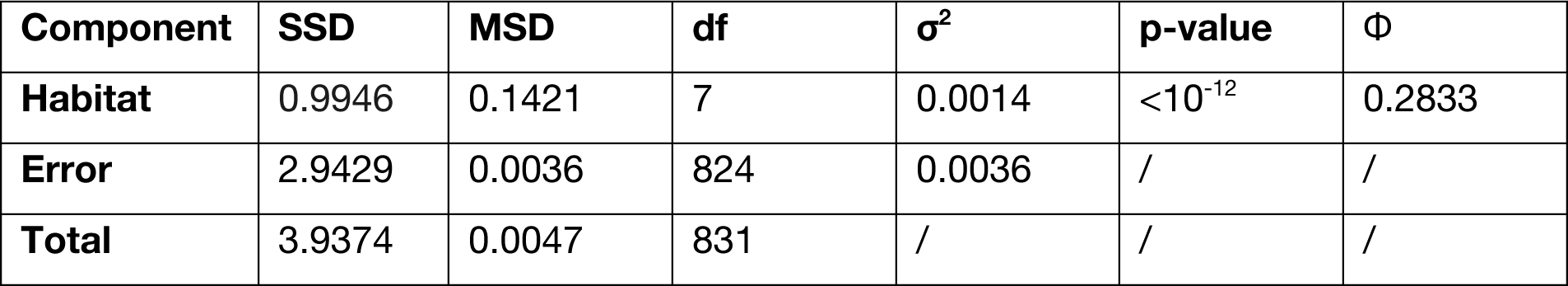
Analysis of molecular variance of genotypes sampled in Ammiad across all years, subdivided by habitat, using only polymorphic sites within Ammiad. Sum of squared deviations (SSD) and mean square deviations (MSD), variance components (σ^2^) and corresponding p-values were computed using the R package *pegas*. The Φ statistic for habitat was used as an estimate of the proportion of molecular variance explained by the habitat structure.

| Component | SSD | MSD | df | $\sigma^2$ | p-value | $\Phi$ |
| --- | --- | --- | --- | --- | --- | --- |
| Habitat | 0.9946 | 0.1421 | 7 | 0.0014 | $<10^{-12}$ | 0.2833 |
| Error | 2.9429 | 0.0036 | 824 | 0.0036 | / | / |
| Total | 3.9374 | 0.0047 | 831 | / | / | / |

**Table S2B.** Analysis of molecular variance of genotypes sampled in Ammiad across all Habitats, subdivided by year, using only polymorphic sites within Ammiad. Sum of squared deviations (SSD) and mean square deviations (MSD), variance components (σ^2^) and corresponding p-values were computed using the R package *pegas*. The Φ statistic for year was used as an estimate of the proportion of molecular variance explained by the temporal structure.

| Component | SSD | MSD | df | $\sigma^2$ | p-value | $\Phi$ |
| --- | --- | --- | --- | --- | --- | --- |
| Year | 0.1197 | 0.0150 | 8 | 2.8618e-06 | 0.4226 | 0.00019 |
| Error | 12.1003 | 0.0147 | 823 | 1.4703e-02 | / | / |
| Total | 12.2200 | 0.0047 | 831 | / | / | / |

**Table S2C.** Analysis of molecular variance of genotypes sampled in Ammiad, modelling both Habitat and time, using only polymorphic sites within Ammiad. Sum of squared deviations (SSD) and mean square deviations (MSD), variance components (σ^2^) and corresponding p-values were computed using the R package *pegas*. Year was modelled as nested within habitat. Note that the low variance explained by time is not due to this (Table S2B). The Φ statistic for habitat and year was used as an estimate of the proportion of molecular variance explained by the year and habitat structure.

| Component | SSD | MSD | df | $\sigma^2$ | p-value | $\Phi$ |
| --- | --- | --- | --- | --- | --- | --- |
| Habitat | 3.1348 | 0.4478 | 7 | 4.3370e-03 | <10<sup>-12</sup> | 0.2831 |
| Year | 0.7146 | 0.0112 | 64 | 1.3254e-05 | 0.7902 | 0.0012 |
| Error | 8.3706 | 0.0110 | 824 | 1.1014e-02 | / | / |
| Total | 12.2200 | 0.0147 | 831 | / | / | / |

**Table S3.** Linear discriminant analysis dimension values.

|  | LD1 | LD2 | LD3 | LD4 | LD5 | LD6 | LD7 |
| --- | --- | --- | --- | --- | --- | --- | --- |
| Anthesis date | -0.005 | -0.528 | -0.346 | 0.177 | -0.198 | -0.053 | 0.124 |
| Coleoptile colour | -0.576 | -0.524 | -0.194 | -0.525 | 0.285 | -0.904 | -0.779 |
| Erect growth habit | -0.822 | 0.32 | 0.746 | -0.985 | 0.188 | 0.336 | -1.242 |
| Germination date | 0.168 | 0.398 | -0.067 | -0.111 | 0.533 | -0.148 | 0.24 |
| Main tiller flag leaf width | 0.739 | -0.27 | -0.828 | -0.49 | -0.28 | -0.355 | 0.21 |
| Main tiller length | 0.759 | -0.497 | -0.328 | 0.675 | 0.144 | -1.084 | 0.159 |
| Main tiller number of spikelets | -0.036 | 0.054 | 0.002 | 0.106 | 0.08 | 0.16 | 0.02 |
| Number of tillers | 0.056 | -0.096 | -0.052 | -0.044 | 0.105 | -0.012 | 0.231 |
| Plant base pigment during tillering | 0.617 | -0.299 | -0.052 | 0.058 | -0.049 | 0.752 | -0.062 |
| Seed area | -1.083 | -1.866 | -2.63 | 0.055 | -2.035 | -1.184 | 0.422 |
| Seed length | 1.344 | 0.816 | 2.349 | -0.173 | 1.634 | 1.42 | -0.05 |
| Seed width | 1.113 | 1.303 | 1.233 | 1.001 | 1.606 | 1.104 | -0.937 |
| Main tiller spike angle | -0.214 | -0.147 | -0.8 | 0.138 | 0.553 | 0.562 | 0.362 |
| Seed weight | -0.217 | 0.239 | 0.024 | -0.68 | -0.724 | -0.234 | 0.704 |

